# Single-AAV CRISPR editing of skeletal muscle in non-human primates with NanoCas, an ultracompact nuclease

**DOI:** 10.1101/2025.01.29.635576

**Authors:** Benjamin J. Rauch, Aaron DeLoughery, Renan Sper, Sean Chen, Sophia Yunanda, Megan Masnaghetti, Ning Chai, Jason Chen Lin, Alexander Neckelmann, Ymer Bjornson, David Paez Espino, Alyssa Sancio, Christopher Schmitt, Clarissa Scholes, Ria Shah, Pooja Kyatsandra Narendra, Sara Ansaloni, SiuSze Tan, Subhadra Jayaraman Rukmini, Sahana Somaiah, Shravanti Suresh, Shiaki Minami, Stepan Tymoshenko, Will Wright, Siming Xu, James Broughton, Matan Drory Retwitzer, Maggie Bobbin, Dave Yuan, Keith Abe, Mark DeWitt, Bohong Zhang, Lucas B. Harrington

## Abstract

CRISPR gene editing is a transformative technology for addressing genetic diseases, but delivery constraints have largely limited its therapeutic applications to liver-targeted and *ex vivo* therapies. Here, we present the discovery and engineering of NanoCas, an ultracompact CRISPR nuclease capable of extending CRISPR’s reach *in vivo* beyond liver targets. We experimentally screened 176 ultracompact CRISPR systems found in metagenomic data and applied protein engineering approaches to enhance the editing efficiency of NanoCas. The optimized NanoCas exhibits potent editing capabilities across various cell systems and tissues *in vivo* when administered via adeno-associated viral (AAV) vectors. This is accomplished despite NanoCas being approximately one-third the size of conventional CRISPR nucleases. In proof-of-concept experiments, we observed robust editing with our optimized NanoCas in mouse models targeting Pcsk9, a gene involved in cholesterol regulation, and targeting exon splice sites in dystrophin to address Duchenne muscular dystrophy (DMD) mutations. We further tested the efficacy of our NanoCas system *in vivo* in non-human primates (NHPs) resulting in editing levels above 30% in muscle tissues. The compact size of NanoCas, in combination with robust nuclease editing, opens the door for single-AAV editing of non-liver tissues *in vivo*, including the use of newer editing modalities such as reverse transcriptase (RT) editing, base editing, and epigenetic editing.

## Introduction

CRISPR-Cas systems have revolutionized the field of genetics, offering unprecedented precision in gene editing and presenting viable strategies for correcting genetic diseases. Due to their large size, existing CRISPR-Cas systems, including Cas9 (∼1,368 amino acids) and Cas12a (∼1,307 amino acids), while effective, exceed the packaging capacity of single adeno-associated viruses (AAVs), the most widely-used vectors for gene delivery *in vivo*^1^ (Fig. 1a, Extended Data Fig. 1b). As a result, CRISPR gene editing’s clinical applications have been constrained to ex vivo therapies and liver-directed therapies where lipid nanoparticles (LNPs) can be used. A compact, potent CRISPR system able to comfortably fit within the AAV genome would overcome these constraints, enabling specific in vivo gene editing of diverse tissues including the central nervous system, skeletal and cardiac muscle, which are challenging to edit by other methods^2^.

**Figure 1.**
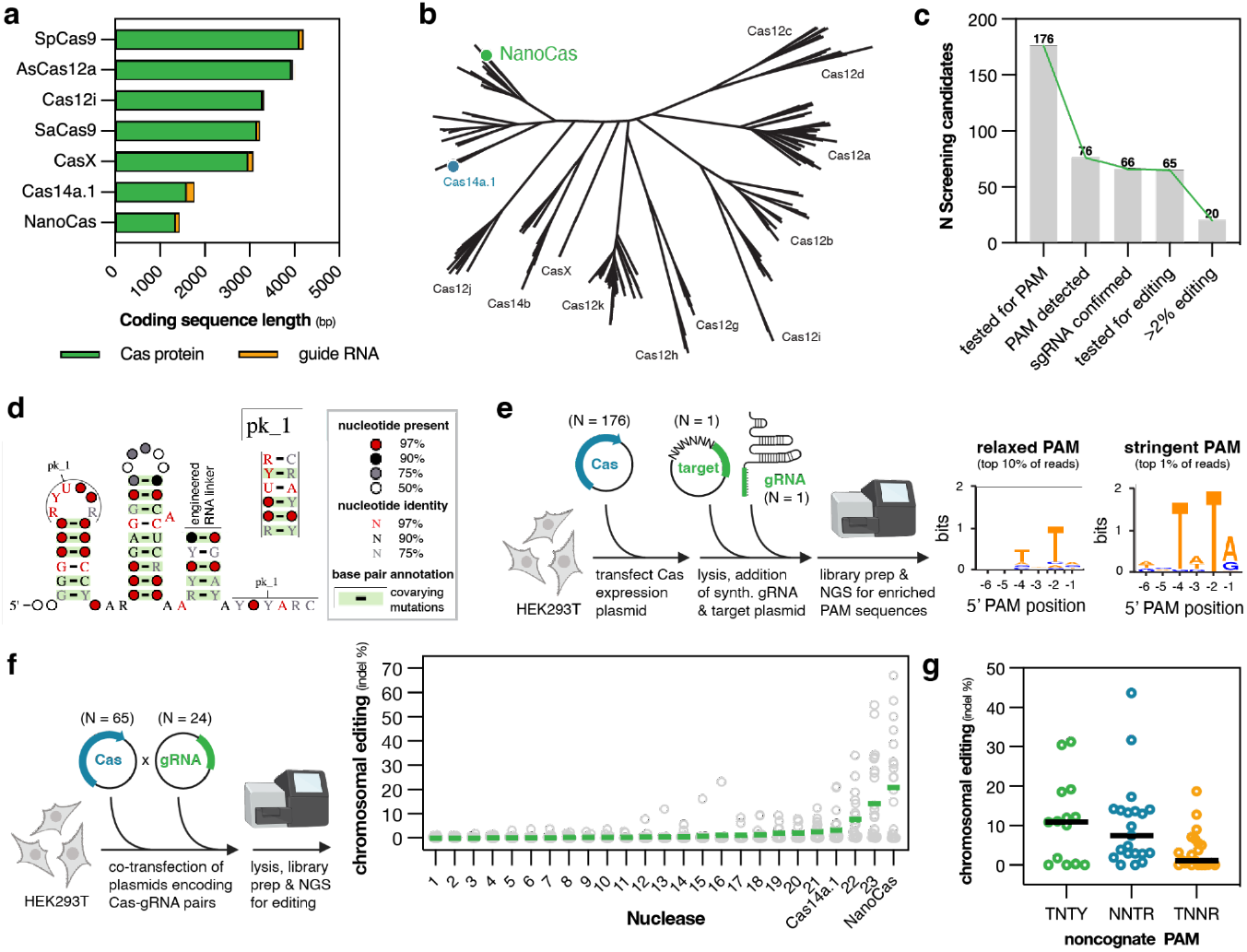
Identification and characterization of NanoCas. (**a**) Coding sequence lengths for Cas nucleases and their corresponding guide RNAs. (**b**) Phylogenetic analysis of Type V CRISPR-Cas nucleases, with NanoCas highlighted in green. (**c**) Screening cascade for NanoCas discovery. (**d**) Consensus guide RNA structure showing pseudoknot (pk_1) and engineered linkers. Nucleotide conservation and base-pairing patterns are indicated per legend. (**e**) PAM identification workflow in HEK293T lysates, with resulting sequence logos shown for relaxed and stringent analyses. (**f**) Chromosomal editing assessment in HEK293T cells. Green lines indicate mean indel frequencies across 24 loci (n=24); individual events shown as grey circles. (**g**) Editing efficiency across targets with non-TNTR PAM sequences.

After the establishment of Cas9 in 2012, extensive efforts have focused on exploring microbial CRISPR diversity and characterizing alternative genome editing platforms (Fig. 1b)^3–7^. Recent analyses suggest that natural diversity encompasses hundreds of thousands of CRISPR systems^8^. The identification of smaller Cas proteins, such as CasX, Cas12i, and SaCas9, has partially addressed the size limitations inherent to SpCas9^6,9,10^. However, even these compact variants remain too large for efficient AAV delivery when combined with necessary regulatory elements and fusion proteins (Fig. 1a). While dual AAV delivery approaches have been attempted to overcome size limitations, these strategies suffer from significantly reduced editing efficiency and create additional manufacturing and safety challenges due to the need to co-deliver multiple vectors. This size constraint particularly impacts newer editing modalities that rely on fused effector proteins to edit DNA, including reverse transcriptase editing^11^, base editing^12^, and epigenetic editing^13^.

The smallest known CRISPR systems to date include Cas14 (Cas12f) and ancestral TnpB, and IscB proteins^14–19^. While promising, these systems demonstrate lower editing efficiency compared to larger systems. Their therapeutic potential remains largely theoretical, with limited *in vivo* studies and no published demonstrations of their efficacy in large animal models such as non-human primates (NHPs). Successful development of new gene editing therapeutics requires robust validation in NHPs, as their high genetic homology with humans and similar tissue architecture and immune responses make them the gold standard for predicting clinical translatability^20^. Furthermore, these systems face additional challenges, including complex protospacer adjacent motif (PAM) sequences, inconsistent guide RNA performance, and lengthy guide RNA sequences, which restrict their utility in therapeutic applications.

These limitations in current CRISPR technology led us to hypothesize that the vast diversity of natural CRISPR systems, combined with advances in protein and RNA engineering, could yield an ultracompact CRISPR system capable of efficient editing within a single AAV vector. We developed a comprehensive screening strategy that simultaneously evaluated multiple parameters critical for therapeutic editing, including size, editing efficiency, specificity, and compatibility with diverse target sequences (Extended Data Fig. 1a). Through this approach, we conducted one of the most extensive screenings of small CRISPR systems to date, synthesizing and experimentally analyzing 176 natural candidate editors to identify and engineer our leading candidate, NanoCas.

NanoCas, with a protein size of approximately 450 amino acids, is roughly one-third the length of SpCas9 and Cas12a and half the size of previously described compact systems like Cas12i, CasX, or SaCas9 (Fig. 1a, Extended Data Fig. 1b). Through our nuclease discovery and engineering efforts, we have demonstrated that NanoCas achieves robust editing efficiency despite its small size. Here we establish NanoCas as an effective tool for *in vivo* genome editing, including through successful editing in non-human primates using a single AAV that targets muscle cells. This work expands the therapeutic possibilities of *in vivo* CRISPR beyond liver-targeted applications, enabling efficient gene editing of previously difficult to reach tissues, and ultimately opens a route to treatment of a number of genetic disorders not addressable with current technologies.

## Results

### Discovery of NanoCas among a cohort of ultracompact CRISPR systems

We began our search for novel CRISPR systems by establishing a Target Editor Profile (TEP) that defined key parameters for therapeutic applications (Extended Data Fig. 1a). The ideal system would: maintain Cas9-equivalent editing efficiency, be smaller than 600 amino acids, target >10% of genomic sequences through PAM recognition, achieve successful editing with >30% of designed gRNAs, match SpCas9’s fidelity, and use an RNA component under 100 nucleotides. Using this TEP as our guide, we conducted a systematic search of 21,980 metagenomes using Hidden Markov models (HMMs) built from known compact Type V CRISPR systems. This initial screen identified 6,945 candidates under 600 amino acids, of which 3,958 contained at least one CRISPR array within 2 kilobases of the nuclease gene.

We implemented a three-stage screening strategy to evaluate these candidates: (1) computational RNA structure prediction, (2) PAM identification and RNA component validation in mammalian cell lysates, and (3) assessment of chromosomal editing in mammalian cells (Fig. 1c).

Our initial focus was on proteins related to Cas14a.1. Based on similarities to Cas14a systems, we hypothesized these candidates would require both a CRISPR-associated RNA (crRNA) and trans-activating RNA (tracrRNA) for function (Fig. 1b)^12^. We identified potential tracrRNAs by analyzing conserved noncoding sequences near cas genes and CRISPR arrays. As expected, our search revealed a diversity of tracrRNA architectures across different candidate clusters. However, in some cases, disparate candidate clusters shared a conserved tracrRNA architecture featuring two stem loops and two unpaired segments with sequence complementary to nearby CRISPR repeats, all within ∼50 nucleotides (Fig. 1d, Extended Data Fig. 1c). This architecture resembled that of Cas14a-like systems but was significantly more compact. We leveraged this finding to conduct RNA-centric database searches, identifying additional candidates that would have been missed through protein-based searches alone. We designed chimeric guide RNAs for all candidates by joining the tracrRNA to target sequences via a 4-nucleotide linker (Fig. 1d), ultimately testing 176 systems—more than half of which employed the ∼50 nt tracrRNA architecture described above.

To evaluate PAM preferences and validate gRNA designs, we expressed candidate nucleases in HEK293T cells rather than using bacterial or in vitro systems. This approach offered the advantage of selecting for variants that express well in mammalian cells—a common translational bottleneck ^21^. We combined cell lysates containing expressed nucleases with synthetic gRNAs and a target plasmid library containing randomized PAM sequences (Fig. 1e). Using NGS analysis of cleaved plasmids^17^, we identified PAM sequences for 76 candidates (43% of those screened, Fig. 1c), revealing approximately 30 distinct PAM preferences.

We then developed a chromosomal editing screen in HEK293T cells to identify nucleases active in human cells and assess their consistency across different target sites (Fig. 2f). As many CRISPR systems show inconsistent performance across gRNA sequences, limiting their utility, we tested each candidate with up to 24 unique gRNAs targeting various genes to comprehensively evaluate editing capabilities. Of the 65 systems tested, 20 showed detectable activity with at least one gRNA, though most achieved average editing rates below 5%. One candidate, which we named NanoCas, distinguished itself with superior performance— achieving an average editing rate of ∼20% and successfully editing with 60% of tested gRNAs. This substantially exceeded other candidates, including our Cas14a.1 control which averaged 8% editing efficiency.

**Figure 2.**
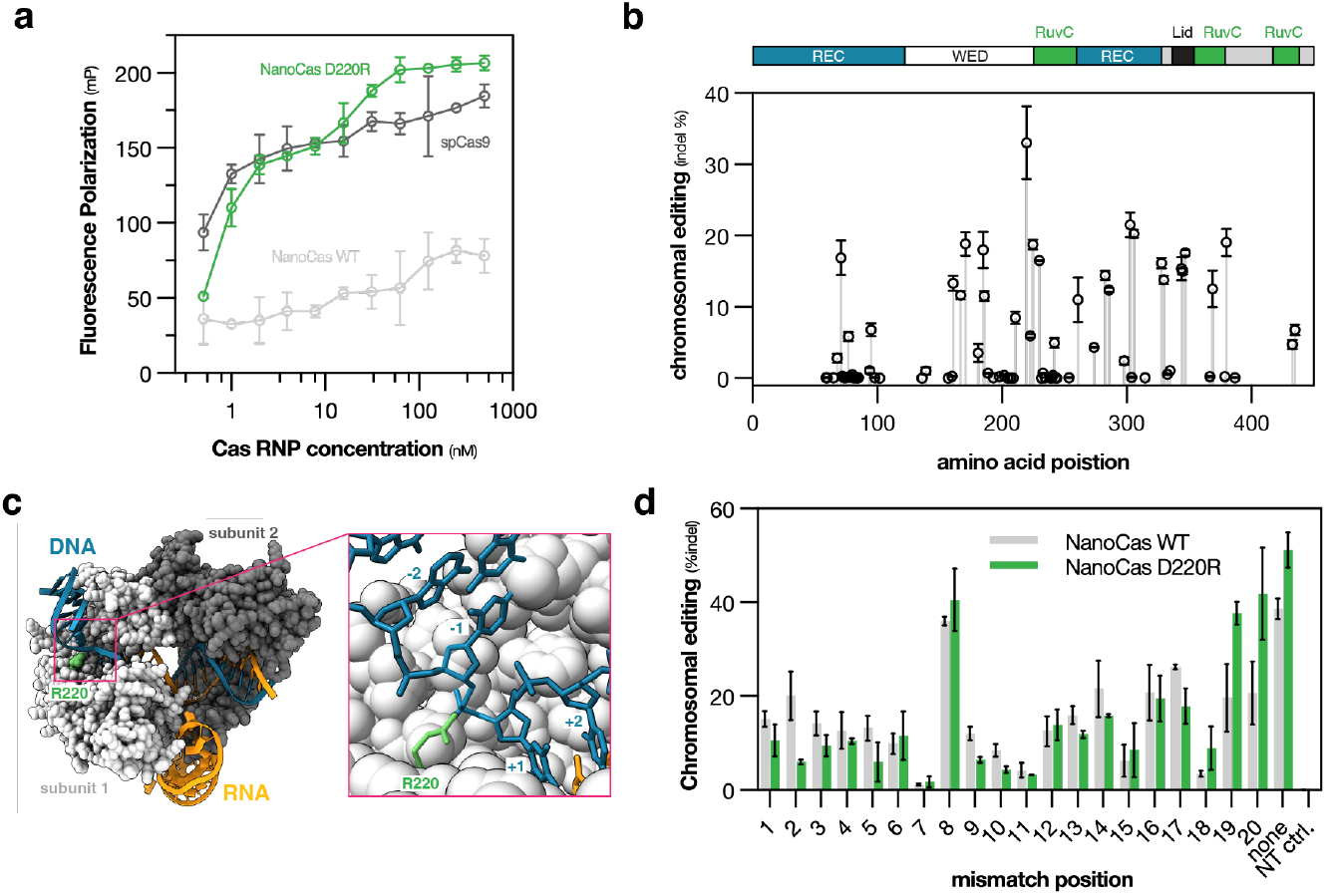
Engineering of an enhanced NanoCas genome editor. (**a**) DNA binding affinity measured by fluorescence polarization for catalytically inactive NanoCas variants and spCas9. (**b**) Chromosomal editing efficiency of NanoCas variants with arginine substitutions, assessed at sub-saturating plasmid doses in HEK293T cells. Domain architecture shown above. (**c**) Structural model of NanoCas D220R-guide RNA-target DNA complex. Inset shows R220 side chain (green) in close proximity to the phosphate group bridging the last PAM (-1) and the first target (+1) nucleotides on the strand that hybridizes to the guide RNA proximally to the PAM-target junction phosphate. (e) Mismatch tolerance analysis comparing wild-type and D220R NanoCas editing at the B2M locus in T cells. Guide RNA mutations were systematically introduced across the spacer sequence.

Initial assessment in cell lysate revealed NanoCas recognized a TNTR PAM sequence (or TNTN under relaxed conditions; Fig. 1e). Next, we further scrutinized the PAM of NanoCas in human cells to determine its identity in a chromosomal editing context as opposed to the plasmid-cleavage assay. NanoCas was screened with 79 guides in total, all with non-TNTR PAMs, and we were surprised to find substantial flexibility at the -4 and -1 positions, with some spacers showing tolerance of N and Y identities at these positions, respectively (Fig. 1g). With the implied NNTN PAM flexibility, NanoCas can natively target 48.7% of bases in the human genome, substantially improving targetability relative to previous compact type V systems^17^.

Like other Type V nucleases, NanoCas generates large deletions distal to the PAM site (Extended Data Fig. 1d). Sanger sequencing mapped precise cut sites to positions +22/+23 on the target strand and +27/+28/+29 on the non-target strand relative to the PAM (Extended Data Fig. 1e).

This comprehensive screening process, from bioinformatic prediction through rigorous functional testing, highlights the rarity of finding highly efficient nucleases in nature. Of thousands of candidates meeting size requirements, only NanoCas emerged with both compact size and robust editing capability (Fig. 1c).

### An engineered NanoCas variant with increased potency

To enhance NanoCas’s genome editing capabilities, we first investigated its biochemical properties to identify potential limitations. Fluorescence polarization assays using a nuclease-deficient variant revealed notably weak binding affinity to target DNA sequences (Fig. 2a). Based on this finding, we engineered variants with positively charged arginine substitutions across the protein to improve DNA binding affinity. Screening these variants in HEK293T cells identified D220R as the most effective mutation (Fig. 2b). This enhanced variant demonstrated improved editing efficiency across a range of mRNA doses in HEK293T cells (Extended Data Fig. 2a). Fluorescence polarization assays confirmed the D220R variant’s enhanced DNA binding (Fig. 2a), supporting our hypothesis that wild-type NanoCas’s editing efficiency was limited by DNA binding affinity.

To understand the molecular basis for D220R’s improved performance, we generated a structural model of NanoCas D220R bound to target DNA (Fig. 2c). The model places R220 near the target strand’s phosphate backbone at the PAM-target interface, suggesting it enhances DNA binding through sequence-independent interactions. This predicted structure aligns with known architectures of Cas12f systems^22–24^, showing two NanoCas protomers bound to a single gRNA-DNA complex.

Given D220R’s enhanced DNA binding, we investigated its impact on editing specificity. We assessed mismatch tolerance in T cells using a guide RNA targeting the B2M locus. By introducing single mismatches at each position across the guide sequence, we found that both wild-type NanoCas and D220R maintain similar mismatch sensitivity profiles, with positions 8-9 and positions 18-20 showing higher tolerance for mismatches (Fig. 2d, Extended Data Figs. 2b). For both variants, mismatch tolerance was particularly pronounced near the PAM-distal end, where editing rates reached approximately 40-50% with D220R.

### NanoCas editing in primary cells and *in vivo* mouse editing via single AAV in the liver

To evaluate NanoCas’s therapeutic potential *ex vivo*, we generated NanoCas mRNA and confirmed the impact of the D220R mutation through dose-response studies in both T cells (TRAC) and CD34+ HSPCs (BCL11A) (Fig. 3a). The engineered variant demonstrated superior editing across all mRNA doses tested, reaching ∼90% efficiency in T cells and >80% in HSPCs, compared to wild-type’s lower maximal efficiencies of ∼80% and ∼50% respectively. This enhanced performance was particularly evident at lower mRNA doses in both cell types.

**Figure 3.**
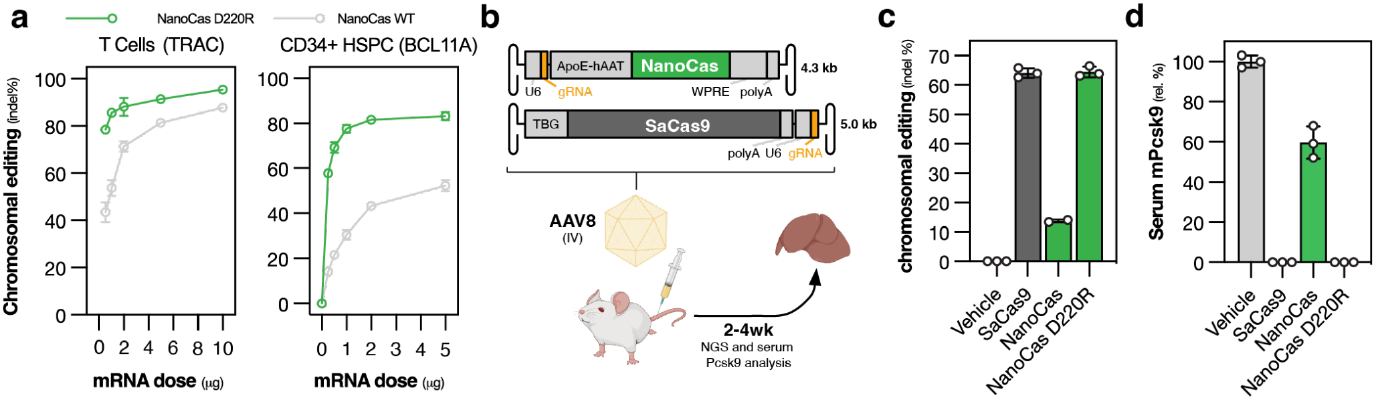
In vivo genome editing efficiency of NanoCas in mouse liver. (**a**) Dose-dependent editing by wild-type and D220R NanoCas delivered as mRNA via nucleofection to human T cells (TRAC locus) and CD34+ HSPCs (BCL11A enhancer). (**b**) Experimental design for comparing NanoCas and SaCas9-mediated Pcsk9 editing via delivery of 2e13vg/kg of AAV8 via tail vein injection. (**c**) Pcsk9 editing efficiency two weeks post-treatment. (**c**) Corresponding serum PCSK9 protein levels.

To evaluate NanoCas’s therapeutic potential *in vivo*, we targeted Pcsk9, a clinically validated gene target for cholesterol reduction. We first screened 114 guide RNAs targeting Pcsk9 exons through plasmid transfection in Hepa1-6 cells (Extended Data Fig. 2c). After identifying the most effective guide RNA, we constructed AAV8 vectors for delivery in wild-type C57BL/6 mice. For SaCas9, we based our AAV on previously used promoters selecting a minimal TBG promoter ^25^. For NanoCas, its smaller size enabled us to include additional regulatory elements while maintaining a total cargo size of just 4.3 kb, well under the AAV packaging limit. We leveraged this extra capacity to incorporate both a larger liver-specific ApoE-hAAT promoter and a Woodchuck Hepatitis Virus Posttranscriptional Regulatory Element (WPRE), which has been shown to enhance mRNA stability^26^ (Fig. 3b).

Following tail vein administration of 2e13 viral genomes/kg, we assessed liver editing efficiency after two weeks. Amplicon sequencing of liver tissue revealed that the NanoCas D220R variant achieved robust editing rates of ∼60%, comparable to SaCas9, while wild-type NanoCas reached 15% editing (Fig. 3c). The 60% editing efficiency suggests successful modification of the majority of hepatocytes^27^. Importantly, both SaCas9 and NanoCas D220R treatments reduced serum PCSK9 protein to undetectable levels, while wild-type NanoCas achieved a 40% reduction (Fig. 3d). These results demonstrate that our engineered NanoCas variant matches the effectiveness of the larger SaCas9 system while maintaining the advantages of a more compact size.

### NanoCas editing of humanized dystrophin in mice

Following our success with liver editing, we explored NanoCas’s potential for therapeutic genome editing in muscle tissue, focusing on dystrophin, which is mutated in Duchenne muscular dystrophy (DMD). Previously described DMD editing strategies include (1) disrupting splice acceptor sites and (2) directly disrupting and reframing premature stop codons and these strategies represent a promising approach for treating DMD by producing truncated but partially functional dystrophin proteins^28^.

To evaluate NanoCas’s capability for exon skipping, we first screened guide RNAs targeting splice acceptor sites in the dystrophin gene using HEK293T cells (Fig. 4a). We identified two promising guides targeting exons 44 and 51, respectively. We packaged these guides along with NanoCas into AAV9-4A vectors, a capsid variant optimized for muscle delivery^29^, using the muscle-specific Ck8e promoter ^30^(Fig. 4b). Initial testing in human myotubes demonstrated robust editing efficiency, with NanoCas achieving 20-40% editing across a range of viral doses (MOI 1e4-1e6), while SaCas9 editing remained below 15% (Extended Data Fig. 4a).

**Figure 4.**
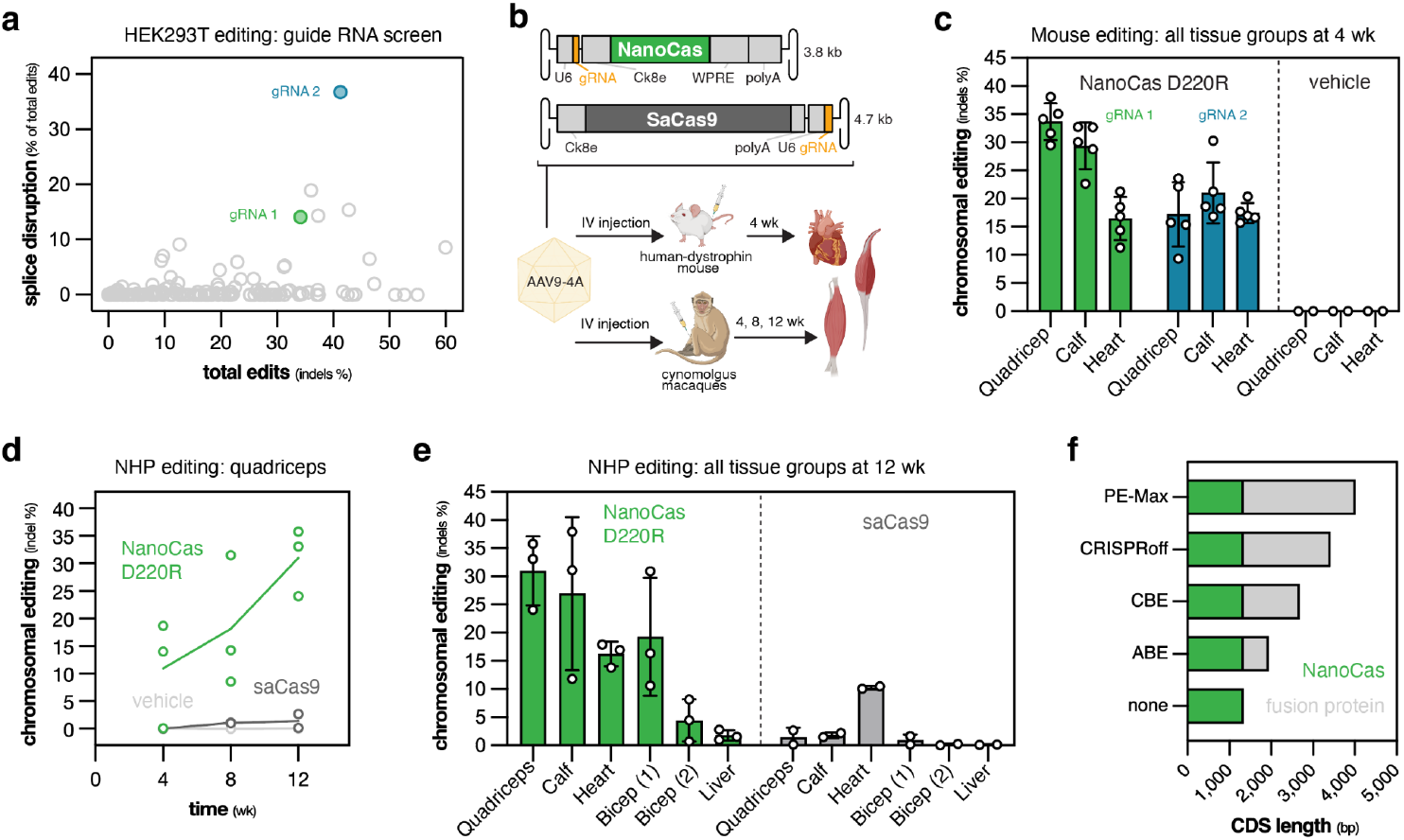
NanoCas-mediated DMD editing in mouse and nonhuman primate models. (a) Guide RNA screen in HEK293T cells showing splice-disrupting versus total editing efficiency. Selected guides (gRNA 1, green; gRNA 2, blue) used for *in vivo* studies. (b) Experimental design for AAV-delivered DMD editing in human dystrophin transgenic mice and cynomolgus macaques. (c) Chromosomal editing of DMD in muscle groups of human-dystrophin mice, 4 weeks post AAV injection. (**e**) Cross-tissue comparison of DMD editing efficiency in NHP at 12 weeks post-treatment, showing editing levels in major muscle groups: quadriceps (vastus lateralis), calf (gastrocnemius), bicep (biceps brachii and biceps femoris) and heart, along with liver. (f) Coding sequence requirements for various precision editors based on NanoCas architecture.

We then evaluated these constructs in a humanized DMD mouse model, administering ∼8e13 of the AAV vectors carrying guide 1 and guide 2, respectively. Analysis of muscle tissue four weeks post-injection revealed substantial editing, ranging from 10-40% across quadriceps, heart, and calf muscles (Fig. 4c). We note that SaCas9 could not be evaluated in this model due to PAM sequence incompatibility with a single nucleotide polymorphism in the human dystrophin sequence in this mouse model.

### NanoCas editing in non-human primates via single AAV

Building on our successful mouse studies, we evaluated NanoCas in NHPs, a critical model for assessing therapeutic gene editing safety and efficacy. We administered our optimized AAV constructs to cynomolgus macaques (n=3) and collected serial muscle biopsies at 4 and 8 weeks, followed by comprehensive tissue analysis at necropsy.

Biopsies of vastus lateralis showed progressive editing, with NanoCas achieving 5-20% indels at 4 weeks and increasing to 10-30% by 8 weeks (Fig. 4d). In contrast, SaCas9-treated animals showed minimal editing (<2%) in the same tissue. At necropsy, we analyzed multiple segments (proximal, medial, and distal) of each muscle tissue to assess editing distribution. Despite some variability between segments, comprehensive analysis of skeletal muscle tissues revealed editing levels up to 30% with NanoCas, while SaCas9 editing remained below 3% (Fig. 4d, 4e, Extended Data Fig. 4b). Notably, these editing levels are particularly encouraging given that restoration of approximately 10% of normal dystrophin levels is thought to be sufficient for therapeutic benefit in DMD patients ^31^.

We also examined editing in cardiac tissue, where NanoCas achieved 15% editing across the left ventricle, left atrium, and septum, compared to 10% with SaCas9 (Fig. 4e). Analysis of liver tissue showed minimal off-target editing, with indel levels below 2% across all four lobes. Throughout the study, animals maintained normal aspartate transferase levels after an initial, transient elevation that resolved by day 8 (data not shown).

To our knowledge, this represents the first demonstration of efficient muscle editing in NHPs using a single-AAV CRISPR system. Previous attempts at in vivo muscle editing in large animals have been limited by the need for dual-vector systems or alternative delivery approaches, highlighting the unique therapeutic potential of NanoCas’s compact size.

## Discussion

Here we present NanoCas, an ultracompact CRISPR nuclease that challenges a fundamental assumption in genome editing: that smaller systems must sacrifice performance for size.

Discovering this system required extensive screening of naturally occurring CRISPR variants - evaluating over 150 candidates in the lab - followed by targeted protein engineering to enhance editing efficiency. The result is a nuclease that matches the capabilities of conventional systems at less than half their size.

The discovery of NanoCas provides insights into how evolution can generate unexpected solutions to complex molecular challenges. While Class 2 CRISPR effectors were thought to require large protein architectures to support their molecular machinery^5^, diverse microbial environments - including nanoarchaea, candidate phyla radiation, and viruses^7^’^16^’^17^ - have driven the development of remarkably compact systems. NanoCas and other compact tsype V systems elegant solution of forming a homodimer on a single guide RNA represents an architectural innovation that would have been nearly impossible to achieve through traditional protein engineering approaches like those used to miniaturize Cas9^18^.

Our systematic characterization of 150 related CRISPR systems revealed the tremendous diversity of CRISPR activity in mammalian cells. While previous screens of Cas12a, a type V CRISPR system, found significant activity in human cells^20^, we observed substantially lower activity rates among ultracompact systems. The reasons for this difference remain unclear but may involve variations in molecular architecture or cellular interactions. Nevertheless, NanoCas demonstrates that carefully selected compact systems can achieve robust editing across various contexts, challenging the assumption that small CRISPR systems are inherently less effective.

We have demonstrated NanoCas’s therapeutic potential through both in vitro studies and in vivo applications in disease-relevant contexts. In the liver, we achieved efficient PCSK9 knockdown in mice. More significantly, we successfully edited the DMD gene in both mouse and non-human primate muscle tissue using a single AAV vector, presenting a potential therapeutic approach for DMD patients who currently have limited treatment options. The robust indel formation at splice junctions in these animal models is particularly promising, and NanoCas’s success in large animal studies is notable given that few CRISPR systems have demonstrated such efficacy.

NanoCas’s compact size opens several promising avenues for therapeutic development. Perhaps most exciting is its potential compatibility with non-double-strand break applications like base editing, reverse transcriptase editing, and epigenetic modification. Even these larger editing modalities can be packaged into a single AAV when built on the NanoCas scaffold (Fig. 4f). Furthermore, NanoCas’s small size - requiring only 2.4kb including regulatory elements - enables the use of self-complementary AAV vectors, which have shown improvements in transduction efficiency^21^’^22^. This could potentially allow for lower dosing requirements, improving the therapeutic window and reducing manufacturing challenges.

Beyond AAV delivery, NanoCas offers advantages for other therapeutic modalities. Its compact size means shorter mRNA constructs for non-viral delivery, potentially improving manufacturing yields and enabling more efficient packaging in lipid nanoparticles (LNPs)^23^. As new delivery technologies emerge, many with their own size constraints^32^, NanoCas’s compact architecture may prove increasingly valuable. This versatility could help extend gene editing therapies to diseases and tissues previously considered inaccessible to conventional CRISPR systems.

The rapid evolution of CRISPR technology from basic research tool to approved therapy in just a decade demonstrates the field’s extraordinary potential. By expanding the capabilities of genome editing while simplifying delivery, systems like NanoCas represent a crucial step toward realizing this potential across a broader range of genetic diseases.

## Acknowledgments

This work is dedicated to the memory of Ning Chai, our dear colleague and friend whose contributions to viral delivery and unwavering commitment to scientific excellence continue to inspire us.

We also thank Janice Chen, Gabor Veres, Emel Alpay, Sara Ansaloni, Sean Coakley, Nick Fantin, Wiputra Hartno, Rachel Herder, Darren Lo, Eric Lu, Jackie Laurel, Bridget McKay, Puloma Sen, Shreyans Sadangi, and Zach Wilson for their guidance and technical assistance.

## Extended Figures

**Extended Data Fig 1.**
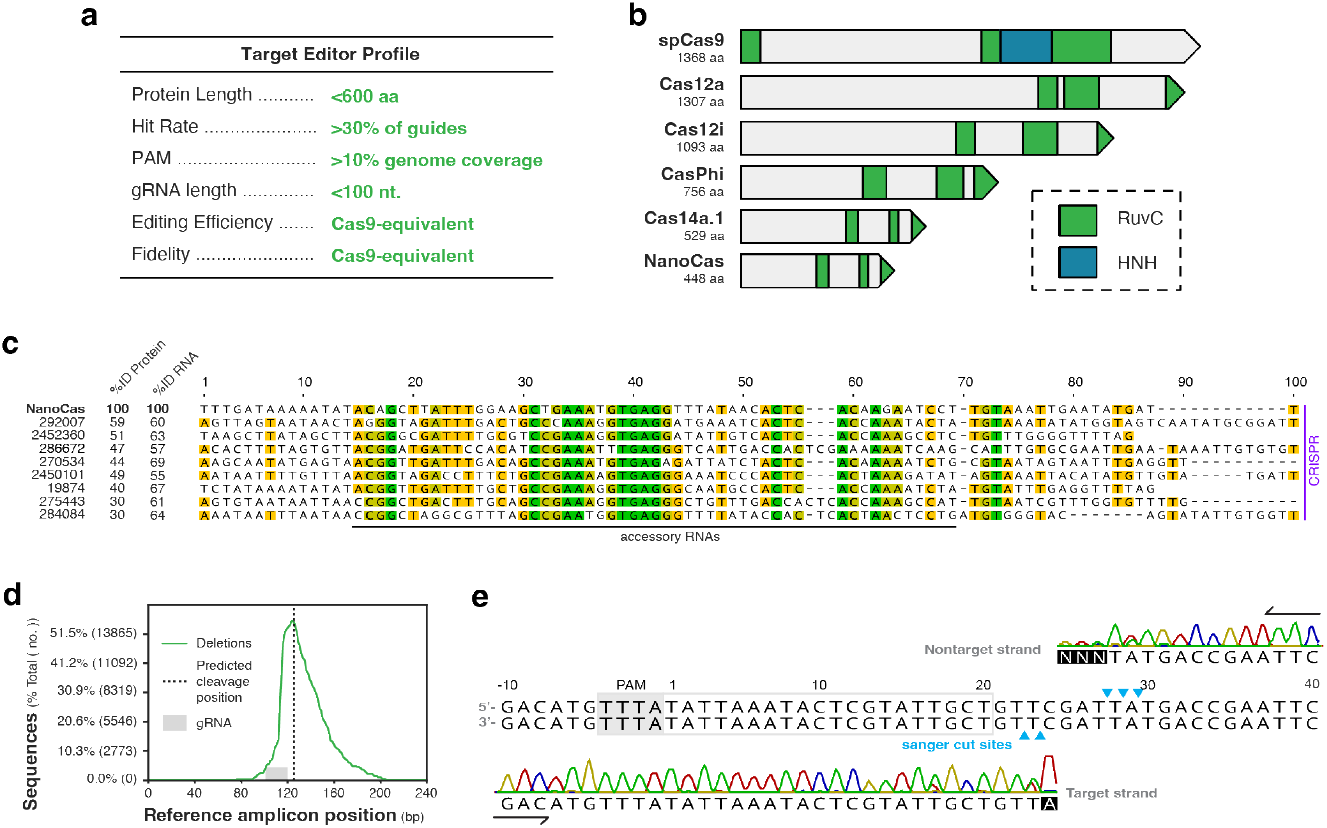
Design criteria and molecular characterization of NanoCas. (a) Design specifications for an ultracompact CRISPR system. (b) Domain architecture of CRISPR-Cas proteins highlighting RuvC and HNH nuclease domains. (c) Conservation analysis of metagenomic sequences upstream of CRISPR arrays associated with NanoCas homologs revealing a highly conserved accessory RNA less than 100 bp upstream from the CRISPR array. Sequence conservation is indicated by color intensity (green: 100%, yellow-green: 80%, yellow: 60%). Purple bar denotes CRISPR array proximity (<50 bp). Percentages show sequence identity to NanoCas protein and RNA. (d) Deletion profile from NanoCas-mediated chromosomal targeting in HEK293T cells, showing preferential PAM-distal cleavage. (e) Cut site mapping via Sanger sequencing of 5’ ends following cleavage of a plasmid by NanoCas. Blue arrows indicate inferred cleavage positions relative to PAM sequence.

**Extended Data Fig 2.**
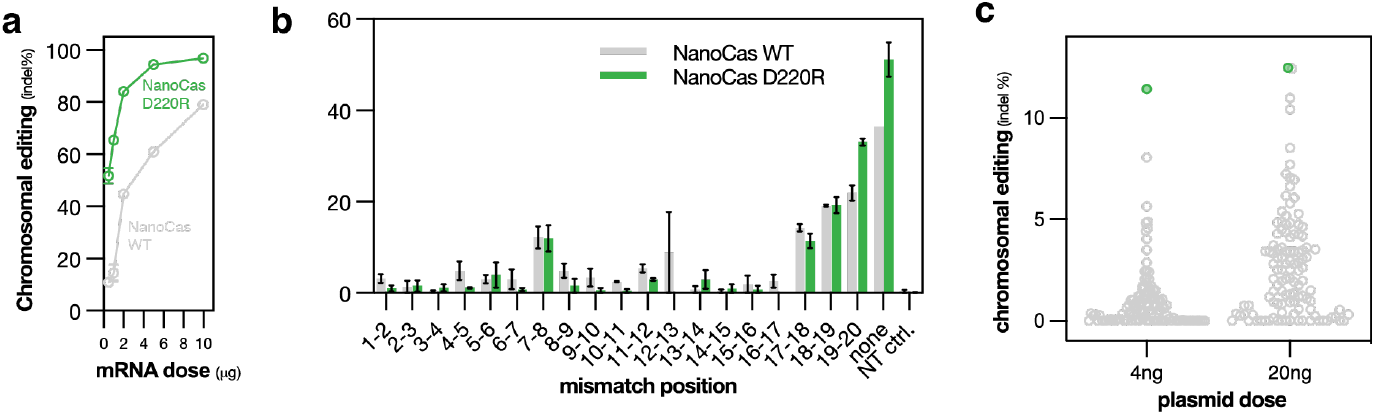
Characterization of enhanced NanoCas D220R specificity. (**a**) Dose-dependent editing of the B2M locus in T cells comparing wild-type and D220R NanoCas delivered as mRNA via nucleofection. (**b**) Double-mismatch tolerance analysis at the B2M locus, with guide RNA mutations systematically introduced across the spacer sequence. (**c**) Guide RNA screen targeting mouse Pcsk9 in Hepa 1-6 cells at two plasmid doses. The best performing guide RNA (green) was selected for AAV delivery.

**Extended Data Fig 4.**
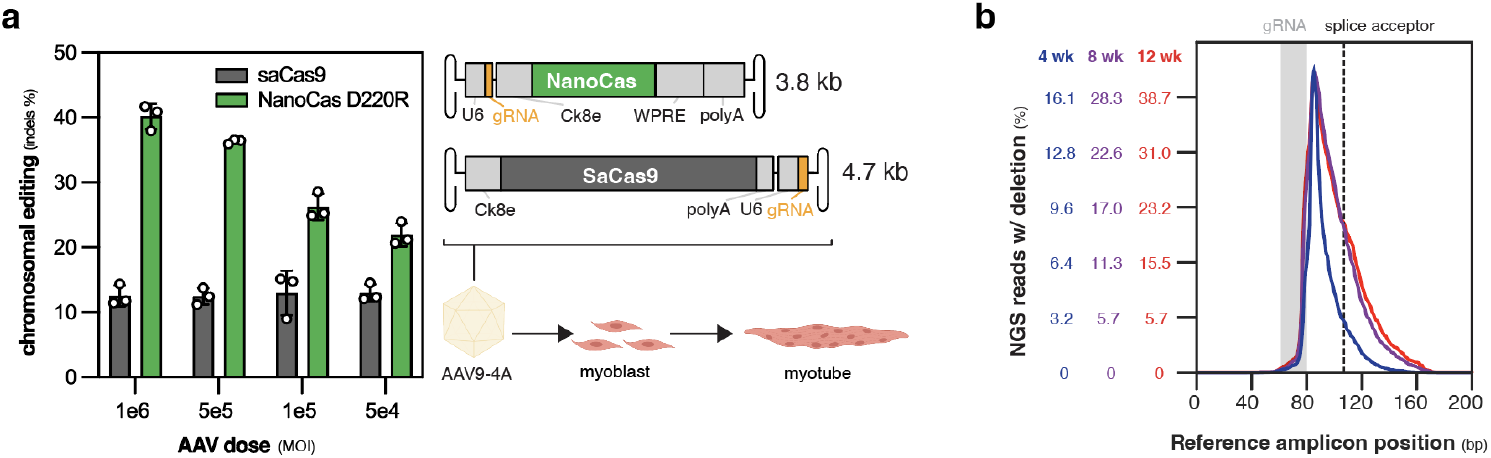
AAV dose response and deletion patterns in DMD editing. (**a**) Dose-dependent DMD editing efficiency in human myotubes *in vitro* comparing NanoCas D220R and SaCas9. AAV delivery and differentiation workflow shown at right. (**b**) Temporal evolution of deletion profiles on dystrophin in NHP quadriceps (vastus lateralis) at 4, 8, and 12 weeks post-treatment, showing progressive enrichment of splice acceptor-disrupting deletions.

## Materials and Methods

### Computational Identification and Analysis of Type V CRISPR Systems

Metagenome sequences (n=21,980) from the JGI Integrated Microbial Genomes and Microbiomes (IMG/M) system^33^ and privately sourced from Darwin Bioprospecting Excellence, S.L. were analyzed using an iterative Hidden Markov Model (HMM) search strategy. Initial HMM profiles were constructed as described previously^34^. Homolog searches were conducted using HMMsearch ^35^ followed by sequence clustering using the MCL algorithm ^36^. Sequences were aligned using Clustal Omega ^37^ and new HMM profiles were generated using hmmbuild.

Sequences were pruned from subsequent iterations if they lacked CRISPR arrays (identified using CRT Tool, ^38^ or auxiliary CRISPR proteins (identified using HMM profiles for known Cas proteins). This process yielded 18,890 type V effector nucleases, which were cataloged with associated CRISPR elements metadata in a custom relational database. Subtype assignment was performed using BLAST searches against a reference database of previously reported type V CRISPR nucleases.

### Genetic Context Analysis and tracrRNA Identification

A clustering-based workflow was implemented to analyze CRISPR locus architecture and identify putative tracrRNAs. Amino acid sequences were clustered using CD-HIT 39 with parameters: 45% minimum identity, word size of 2, and -g 1. For each sequence, 2000 nucleotides flanking the coding sequence were extracted when contig length permitted.

Sequences within clusters of ≥5 members were aligned using MAFFT with default settings. Cluster alignments were analyzed in Geneious with functional annotations including predicted ORFs and CRISPR arrays. For clusters containing nuclease CDS of 1000-1500 bp, conserved non-coding regions were identified and evaluated as tracrRNA candidates using ViennaRNA ^40^. Candidate structures were compared to previously validated tracrRNAs.

### RNA Structure Analysis and Homolog Search

For RNA-centric homolog identification, tracrRNA and repeat sequences from NanoCas-like proteins were concatenated using an AUUU linker. Multiple sequence alignment with structure prediction was performed using LocARNA ^41^, and the linker was replaced with gap symbols. The resulting stockholm format alignment was used to construct a covariance model using Infernal ^42^ for metagenomic database searches. Consensus RNA structural analysis was performed using R2R ^43^.

### PAM Identification

PAM sequences were determined through in vitro enrichment (IVE) using HEK293T-expressed nucleases. Cells were seeded at 30,000 cells/200 µL in 96-well plates and transfected with 300 ng nuclease expression plasmid. After 72 hours, cells were lysed in a buffer containing 1% triton, 150 mM KCl, 20 mM HEPES pH 7.5, 5 mM MgCl2, 1 mM DTT, and 5% glycerol. Nuclease-containing lysate (10 µL) was complexed with guide RNAs (2 µL, 500 nM) for 15 minutes at 37°C. Complexes were added to IVE reaction mix containing 1,000 ng of 5’ PAM library in 1x Cutsmart buffer (100 µL total) and incubated for 60 minutes at 37°C followed by 15 minutes at 45°C. Reactions were terminated with proteinase K (1 µL) and EDTA (5 µL, 500 mM).

Library preparation was performed using NEBNext Ultra II workflow, beginning with End Repair/dA-tailing. Two sequential PCRs were performed: first using IVE_F_A-D primer pool and IVE_R primer (8 cycles), followed by Illumina indexing primers (13 cycles). PCR conditions: 98°C for 30 seconds; cycles of 98°C for 10 seconds, 65°C for 15 seconds; final extension at 65°C for 1 minute. PAM sequences were analyzed from MiSeq 2×150 reads, and sequence logos were generated using WebLogo5.

### Guide RNA Synthesis

Guide RNAs <103 nucleotides were chemically synthesized (Synthego). Longer guides were produced by in vitro transcription (IVT) using PCR-generated templates (gBlock, IDT) and Q5 mastermix (NEB). IVT reactions (100 µL) contained template DNA, T7 polymerase (NEB), 5 mM dNTPs, and Murine RNase Inhibitor (NEB). After 4 hours at 37°C, reactions were DNase I-treated (30 minutes, 37°C), terminated with 0.5 M EDTA, and purified using RNAcleanXP beads (1.8:1 ratio).

### Protein Production and Characterization

NanoCas proteins were expressed with N-terminal His(10)-MBP-TEV tags in NiCo21(DE3) E. coli. Cultures (1L Terrific Broth, 100 µg/mL carbenecillin) were grown at 37°C to OD600 0.6, induced with 0.5 mM IPTG, and expressed overnight at 18°C. Cells were lysed by sonication in protease inhibitor-containing buffer. Proteins were purified sequentially using: (1) HisTrap FF nickel column (2) TEV protease cleavage (3) Tandem MBP-Trap and Heparin HP columns (4)Final polishing on HiLoad Superdex 200 16/600

The final buffer contained 20 mM HEPES, 500 mM NaCl, 10% glycerol, 1 mM TCEP, pH 7.5. Protein purity (>95%) was confirmed by SDS-PAGE and analytical SEC.

### Fluorescence Polarization Binding Assays

RNP complexes were formed by incubating NanoCas protein with gRNA (1 µM each) in 20 mM HEPES, 1 mM TCEP, 0.2 mg/mL BSA for 15 minutes at room temperature. Serial dilutions (0.5-500 nM) were incubated with 0.1 nM 3’ 6-FAM labeled DNA probes in assay buffer (20 mM HEPES pH 7.5, 0.2 mg/mL BSA, 1 mM TCEP, 100 mM NaCl, ±5 mM Magnesium acetate/EDTA). Reactions were measured in black 384-well plates after 30 minutes at 37°C using a Biotek-Synergy H2 plate reader. KD values were determined using non-linear fit curves (one site-specific binding) in PRISM software.

### Cell Culture and Transfection

HEK293T and Hepa1-6 cells (ATCC) were maintained under standard conditions. For editing experiments, HEK293T cells were seeded at 30,000 cells/200 µL in 96-well plates and transfected with up to 300 ng total plasmid DNA using TransIT-293 (Mirus Bio) in Optimem. Hepa1-6 cells were seeded at 20,000 cells/well and transfected using Lipofectamine 3000 following the manufacturer’s protocol. Editing outcomes were assessed after 48-72 hours.

### Primary Cell Editing

CD34+ HSPCs (AllCells) were cultured in SFEM II medium containing 1X CC110 cytokine cocktail and 1% penicillin-streptomycin for 2 days pre-editing. Cells were nucleofected using the Neon system (1500V/10ms/3pulses, Buffer “T”) with NanoCas mRNA and BCL11A-targeting guide RNA (500 pmol). Post-editing, cells were assessed for viability and transferred to erythroid expansion medium for 4 days before analysis.

Human CD3+ Pan T cells (AllCells) were cultured in X-VIVO 15 medium supplemented with 5% Human AB serum, 2mM L-Glutamine, 1% Penicillin-streptomycin, and cytokines (200U/ml IL-2, 10 ng/ml each IL-7 and IL-15). T cells were activated using CD3/CD28 Dynabeads for 72 hours before nucleofection. Editing was performed using Amaxa 4D (pulse code EH115, P3 Primary Cell kit) in 20 µL total volume. For dose-response studies, varying NanoCas mRNA amounts (0.5-10 µg) were combined with 500 pmol guide RNA.

### Guide RNA Specificity Studies

Mismatch tolerance was evaluated by introducing single and double mutations across positions 0-20 of the guide RNA spacer sequence. For TRAC off-target analysis, potential sites were predicted using Cas-Offinder. Both on-target and predicted off-target sites were assessed by NGS analysis.

### AAV Production and Characterization

AAVs were produced in Suspension Viral Production Cells 2.0 using triple transfection (AAV Rep-Cap plasmid, adenovirus helper plasmid, and ITR-flanked transfer plasmid; 1:1:1 ratio) with PEIpro. After 3 days, viral particles were harvested from cells and medium using AAV-Max Lysis Buffer with Benzonase treatment. Purification was performed by sucrose or iodixanol gradient ultracentrifugation. Vector quality was assessed by Titer determination (qPCR), Genome integrity (alkaline gel electrophoresis), Purity (SDS-PAGE), Endotoxin content (<0.5 EU/mL threshold).

### *In vitro* AAV Assessment

Immortalized human myoblasts were transduced in suspension (MOI range: 5e4-1e6) with ROCK inhibitor in serum-free medium. After 6 hours, complete growth medium was added. At 48-72 hours post-transduction, differentiation was initiated using Myocult media. Genomic DNA was collected after 28 days of differentiation for NGS analysis.

### In vivo injections and tissue analysis for mPCSK9 study

All mouse experiments were performed in accordance with protocols approved by the CRADL Institutional Animal Care and Use Committee (IACUC). C57BL/6J mice (JAX #000664), male, 6– 8 weeks of age, were obtained from Jackson Laboratories were and kept on a 12-h light/12-h dark schedule with food and water provided *ad libitum*.

### mPCSk9 ELISA and serum analysis

C57BL/6J mouse whole blood was collected by cardiac puncture and serum was isolated by centrifugation of whole blood in a serum separator tube. mPCSK9 serum protein was measured using the Mouse Proprotein Convertase 9/PCSK9 ELISA Kit (Biotechne R&D Systems). Mouse serum was diluted 200-fold and ELISA was processed following manufacturer provided protocol. Sample absorbance was measured using a microplate reader set to 450nm.

### *In vivo* study in DMD mice model

The JAX:018900 hDMD mice model was selected for this study. This mouse model contains The hDMD transgene in a 2.7 Mbp yeast artificial chromosome (YAC) sequence containing 2 copies of the full-length 2.3 Mbp human dystrophin gene (*DMD*) on each allele. The transgene integration site is on mouse chromosome 5, this mouse does not display DMD-associated phenotype. Homozygous animals were selected for the testing of AAV-NanoCas. Genomic sequencing of the target locus for spacer sequence confirmed the presence of spacer and PAM sequence for NanoCas gRNA 1 and gRNA 2, and the target sequence was not present in the mouse DMD locus. Genomic sequencing analysis of the saCas9 targeting region identified a mismatch within PAM sequences, resulting in excluding the AAV-saCas9 treatment group. Homozygous hDMD mice were injected via tail injection with 7.72 e13 vg/kg (guide 1) and 8.06 e13 vg/kg (guide 2) viral particles and monitored throughout the length of the study. Collection of quadriceps, calf, and heart was performed at 4 weeks post-dosing for NGS analysis.

### *In vivo* study in NHP

The NHP study was performed by protocols approved by the CRADL Institutional Animal Care and Use Committee (IACUC). Male cynomolgus macaques (2 to 4 years of age) were selected for the study and screened for the presence of neutralizing antibodies against AAV9-4A and monitored for serum conversation 2-4 weeks before the study started. NHP animals received Methylprednisolone (20 mg/kg, IM) before AAV treatment. AAV dosing was conducted via 30-minute intravenous (Cephalic vein) infusion at 5ml/Kg and 5 e13 VG/Kg dose. Biceps Brachii and Vastus Lateralis biopsy punches were collected at 4 weeks left side) and 8 weeks (right side). Serum samples were collected at various time points throughout the study for clinical pathology assessment. Necropsy was performed at 12 weeks, quadriceps, calf, heart, biceps, and liver were collected for NGS. Skeletal muscle samples were collected at the proximal, medial, and distal regions of each muscle and expressed as the mean value/animal for each site. Heart samples were collected from the left ventricle, left atrium, and septum, and expressed as the mean value.

### DNA Extraction and Library Preparation

Genomic DNA was isolated using either Wizard SV 96 kit (tissue samples) or Quick Extract solution (cells). Target regions were PCR-amplified and indexed using Illumina Nextera primers. Libraries were pooled and sequenced on Illumina MiSeq or NextSeq platforms.

### Protein Analysis

ELISA measurements were performed according to manufacturer protocols with sample-specific dilution factors. Absorbance readings were taken at 450nm using a standard microplate reader. Standard curves were generated using manufacturer-provided reference standards.

### Statistical Analysis and Data Visualization

Data processing and statistical analyses were performed using GraphPad Prism (v9, GraphPad Software Inc.). Specific statistical tests and significance thresholds are detailed in figure legends. Sequence alignments and analysis were performed using Geneious Prime (v2023.1, Biomatters Ltd.).

Sequencing data was processed using CRISPResso2 to quantify editing outcomes. Indel percentages were calculated as the fraction of reads containing insertions or deletions relative to unedited reference sequences. For multi-site analysis (NHP studies), results from different sampling locations within the same tissue were averaged to generate mean values per animal.

Figures were generated using a combination of: GraphPad Prism (v9) for statistical plots and graphs, Geneious Prime (v2023.1) for sequence alignments and analysis, BioRender.com for schematic diagrams, WebLogo5 for sequence logos.

## References

1. Wang, D., Tai, P. W. L. & Gao, G. Adeno-associated virus vector as a platform for gene therapy delivery. Nat. Rev. Drug Discov. 18, 358–378 (2019).

2. Lino, C. A., Harper, J. C., Carney, J. P. & Timlin, J. A. Delivering CRISPR: a review of the challenges and approaches. Drug Deliv. 25, 1234–1257 (2018).

3. Burstein, D. et al. Major bacterial lineages are essentially devoid of CRISPR-Cas viral defence systems. Nature Communications 7, 10613–10613 (2016).

4. Shmakov, S. et al. Diversity and evolution of class 2 CRISPR–Cas systems. Nature Reviews Microbiology 15, 169–182 (2017).

5. Makarova, K. S. et al. An updated evolutionary classification of CRISPR-Cas systems. Nat Rev Microbiol, 1–15 (2015).

6. Burstein, D. et al. New CRISPR-Cas systems from uncultivated microbes. Nature 542, 237–241 (2017).

7. Shmakov, S. et al. Discovery and Functional Characterization of Diverse Class 2 CRISPR-Cas Systems. Molecular Cell 60, 385–397 (2015).

8. Ruffolo, J. A. et al. Design of highly functional genome editors by modeling the universe of CRISPR-Cas sequences. bioRxiv 2024.04.22.590591 (2024) doi:10.1101/2024.04.22.590591.

9. Garrity, A. J. et al. Functionally diverse type V CRISPR-Cas systems. Science 363, 88–91 (2018).

10. Ran, F. A. et al. In vivo genome editing using Staphylococcus aureus Cas9. Nature 520, 186–191 (2015).

11. Anzalone, A. V. et al. Search-and-replace genome editing without double-strand breaks or donor DNA. Nature 576, 149–157 (2019).

12. Komor, A. C., Kim, Y. B., Packer, M. S., Zuris, J. A. & Liu, D. R. Programmable editing of a target base in genomic DNA without double-stranded DNA cleavage. Nature 533, 420–424 (2016).

13. Nuñez, J. K. et al. Genome-wide programmable transcriptional memory by CRISPR-based epigenome editing. Cell 184, 2503-2519.e17 (2021).

14. Harrington, L. B. et al. Programmed DNA destruction by miniature CRISPR-Cas14 enzymes. Science (2018) doi:10.1126/science.aav4294.

15. Karvelis, T. et al. Transposon-associated TnpB is a programmable RNA-guided DNA endonuclease. Nature 599, 692–696 (2021).

16. Saito, M. et al. Fanzor is a eukaryotic programmable RNA-guided endonuclease. Nature 620, 660–668 (2023).

17. Karvelis, T. et al. PAM recognition by miniature CRISPR–Cas12f nucleases triggers programmable double-stranded DNA target cleavage. Nucleic Acids Res. 48, 5016–5023 (2020).

18. Altae-Tran, H. et al. The widespread IS200/605 transposon family encodes diverse programmable RNA-guided endonucleases. Science 374, 57–65 (2021).

19. Xu, X. et al. Engineered miniature CRISPR-Cas system for mammalian genome regulation and editing. Mol. Cell 81, 4333-4345.e4 (2021).

20. Kang, Y., Chu, C., Wang, F. & Niu, Y. CRISPR/Cas9-mediated genome editing in nonhuman primates. Dis. Model. Mech. 12, dmm039982 (2019).

21. Walton, R. T., Hsu, J. Y., Joung, J. K. & Kleinstiver, B. P. Scalable characterization of the PAM requirements of CRISPR–Cas enzymes using HT-PAMDA. Nat. Protoc. 16, 1511–1547 (2021).

22. Takeda, S. N. et al. Structure of the miniature type V-F CRISPR-Cas effector enzyme. Mol. Cell 81, 558-570.e3 (2021).

23. Xiao, R., Li, Z., Wang, S., Han, R. & Chang, L. Structural basis for substrate recognition and cleavage by the dimerization-dependent CRISPR–Cas12f nuclease. Nucleic Acids Res. 49, 4120–4128 (2021).

24. Wu, T. et al. An engineered hypercompact CRISPR-Cas12f system with boosted gene-editing activity. Nat. Chem. Biol. 19, 1384–1393 (2023).

25. Ran, F. A. et al. In vivo genome editing using Staphylococcus aureus Cas9. Nature (2015) doi:10.1038/nature14299.

26. Donello, J. E., Loeb, J. E. & Hope, T. J. Woodchuck Hepatitis Virus Contains a Tripartite Posttranscriptional Regulatory Element. J. Virol. 72, 5085–5092 (1998).

27. Malarkey, D. E., Johnson, K., Ryan, L., Boorman, G. & Maronpot, R. R. New Insights into Functional Aspects of Liver Morphology. Toxicol. Pathol. 33, 27–34 (2005).

28. Matsuo, M. Duchenne/Becker muscular dystrophy: From molecular diagnosis to gene therapy. Brain and Development 18, 167–172 (1996).

29. Tabebordbar, M. et al. Directed evolution of a family of AAV capsid variants enabling potent muscle-directed gene delivery across species. Cell 184, 4919-4938.e22 (2021).

30. Amoasii, L. et al. Single-cut genome editing restores dystrophin expression in a new mouse model of muscular dystrophy. Sci. Transl. Med. 9, (2017).

31. Hoffman, E. P. et al. Restoring Dystrophin Expression in Duchenne Muscular Dystrophy Muscle Progress in Exon Skipping and Stop Codon Read Through. Am. J. Pathol. 179, 12–22 (2011).

32. Prince, C. et al. A novel functional gene delivery platform based on a commensal human anellovirus demonstrates transduction in multiple tissue types. bioRxiv 2024.03.27.586964 (2024) doi:10.1101/2024.03.27.586964.

33. Chen, I.-M. A. et al. IMG/M v.5.0: an integrated data management and comparative analysis system for microbial genomes and microbiomes. Nucleic Acids Res. 47, D666–D677 (2019).

34. Burstein, D. et al. New CRISPR–Cas systems from uncultivated microbes. Nature 542, 237–241 (2017).

35. Johnson, L. S., Eddy, S. R. & Portugaly, E. Hidden Markov model speed heuristic and iterative HMM search procedure. BMC Bioinform. 11, 431 (2010).

36. Enright, A. J., Dongen, S. V. & Ouzounis, C. A. An efficient algorithm for large-scale detection of protein families. Nucleic Acids Res. 30, 1575–1584 (2002).

37. Sievers, F. & Higgins, D. G. Clustal Omega for making accurate alignments of many protein sequences. Protein Sci. 27, 135–145 (2018).

38. Bland, C. et al. CRISPR Recognition Tool (CRT): a tool for automatic detection of clustered regularly interspaced palindromic repeats. BMC Bioinform. 8, 209 (2007).

39. Fu, L., Niu, B., Zhu, Z., Wu, S. & Li, W. CD-HIT: accelerated for clustering the next-generation sequencing data. Bioinformatics 28, 3150–3152 (2012).

40. Lorenz, R. et al. ViennaRNA Package 2.0. Algorithms Mol. Biol. 6, 26 (2011).

41. Will, S., Joshi, T., Hofacker, I. L., Stadler, P. F. & Backofen, R. LocARNA-P: Accurate boundary prediction and improved detection of structural RNAs. RNA 18, 900–914 (2012).

42. Nawrocki, E. P. & Eddy, S. R. Infernal 1.1: 100-fold faster RNA homology searches. Bioinformatics 29, 2933–2935 (2013).

43. Weinberg, Z. & Breaker, R. R. R2R - software to speed the depiction of aesthetic consensus RNA secondary structures. BMC Bioinform. 12, 3 (2011).

